# Behavior and survival of earthworm (*Eisenia andrei*) to exposure to glyphosate-contaminated weed compost

**DOI:** 10.1101/2024.12.30.630776

**Authors:** Maria Luisa Velazquez Vazquez, Gustavo C. Ortiz-Ceballos, Beatriz Yáñez-Rivera, Angel I. Ortiz-Ceballos

## Abstract

Weeds growing in crops contribute to the maintenance of soil biodiversity; for example, they are a source of organic matter and nutrients for soil invertebrates. However, little is known about the impact of glyphosate-contaminated weeds on the behavior and survival of earthworms. The objective of this study was to evaluate the behavior and survival of the earthworm *Eisenia andrei* upon exposure to a glyphosate-contaminated weed compost. The study employed standardized avoidance and acute tests to assess the repellency and mortality of *E. andrei* earthworms at four doses of the commercial herbicide (l/ha): Control = 0, Low = 2, Medium = 4, and High = 6. The avoidance essay results indicated that earthworms with similar biomass did not avoid glyphosate-contaminated weed composts, with avoidance rates < 80%. However, the acute test revealed that increasing the herbicide dose led to a reduction in earthworm growth and mortality. Consequently, it was concluded that weed composts with glyphosate exerted a toxic effect on the survival of *E. andrei*. It is recommended that future studies focus on the microbiota associated with *E. andrei* in the decomposition of the weed and the degradation of the herbicide.

## Introduction

Earthworms are key organisms in the disintegration of organic matter and are able to help degrade xenobiotics (Zeb et al. 2020; Dada et al. 2021). The process of such activities is mediated in the digestive tract of earthworms by bacterial colonies (Ceceña et al., 2022; Nárvaez et al., 2022), which can be modified by diet, stress, contact with xenobiotics, or disposal of organic matter (Yang et al., 2022; Wang et al., 2023).

The necessity of ecotoxicity tests for the assessment of soil contamination has been underscored by numerous studies (Wang et al. 2016; Carriquiborde et al. 2021). Their importance is such that they are implemented and regulated at both the international and national levels (ISO 2008; ISO 2012; OECD 2016). In Mexico, the International Organization for Standardization (ISO) employs the earthworm of the genus *Eisenia* as a model, given its status as a crucial indicator of soil health and quality (Casabé et al., 2007; Zerbino 2007). This selection is attributed to the earthworm’s physiological characteristics and its capacity to avoid contaminated environments (Falco, 2007; García-Pérez et al., 2014; Palafox et al., 2021).

The ISO 17512-1: 2008 avoidance essay consists of placing an environment with xenobiotic next to a control one, so that earthworms move between them in 48 hours. If 80% of the earthworms are in the control, it indicates that they reject the contaminated environment, otherwise they are indifferent (ISO 2008). The ISO 11268-1:2012 survival essay, on the other hand, consists of exposing earthworms only to the contaminated environment, and evaluates the loss of biomass, epithelial changes and death at 7 and 14 d of exposure; if an earthworm does not move upon contact stimulation, it is considered dead and the percentage of mortality per treatment is evaluated (ISO 2012).

The presence of glyphosate has been identified in crop fields where the herbicide is applied (Salazar et al., 2016; Mamy et al., 2010; Helander et al., 2018), also in surrounding fields where residues are incorporated by leaching and runoff (Gaupp-Bergahusen et al., 2015; González-Chávez, 2019). Herbicide persistence times have been identified to vary depending on environmental conditions (Zirena et al., 2018; Muskus et al., 2020), as well as application rate and frequency (Carriquiborde et al., 2021; Lorch et al., 2021). Glyphosate residues in soil range from 6 to 20 days while in plant matter they reach 55 to 80 days (Doublet et al. 2006; Mamy et al., 2016).

On the other hand, glyphosate has been documented to induce weight reduction in *Eisenia andrei* (Piola et al., 2013) and *E. fetida* (Yasmin and Souza 2007; Correira and Moreira 2010); Furthermore, glyphosate has been observed to disrupt the reproductive activity of *Lombricus terrestris* (Gaupp-Berghausen et al., 2015) and *Pontoscolex corethrurus* (García-Perez et al., 2016).

In the Xalapa-Coatepec region, the coffee agroecosystem is a significant resource that has been impacted by intensive agricultural management practices (Espinoza-Guzmán et al., 2020; Ortiz-Ceballos et al., 2020). The interaction between glyphosate, coffee plant weeds, and soil organisms has not been characterized. Consequently, the objective of this study was to evaluate the effect of glyphosate-contaminated coffee weeds on the behavior and survival of the tropical earthworm *E. andrei* through avoidance and acute test essays.

## Materials and methods Study area

In the locality of Las Lomas, Coatepec, Ver. (19°26’20.5.5’’ N, 96°54’01.1’’ W), a plot with a diversified shade coffee agroecosystem was selected, in whose history no herbicide control of weeds was reported. Within the plot, a 100 m2 quadrat was divided into eight plots (2×4 m), leaving corridors between them (1 m).

### Characterization of weeds

Prior to herbicide application, 16 subplots of 0.5 × 0.5 m were randomly delimited and weeds samples were systematically collected from these subplots for taxonomic classification and assessment of nutritional quality. Additionally, the diversity, abundance, and biomass of the collected weeds were also determined.

### Application and analytical determination of glyphosate

A completely randomized design with four treatments was installed, each treatment consisted of a commercial dose of glyphosate (L/ha): Control = 0, Low = 2, Medium = 4 and High = 6; the hand pump used for the application of the herbicide in the experimental plots was previously calibrated following the technical recommendation (Marchesi and Pauletti, 2023). At 10 days after application (Salazar et al., 2016), the weeds of each treatment were harvested standing; then, they were placed in plastic bags, then dried at ambient temperature, and finally, subsamples were taken for the analytical determination of glyphosate.

### Analytical determination of glyphosate

In the laboratories of the Food and Development Center A.C (Culiacán Campus), the glyphosate content in weeds was determined based on the protocol Rapid method for the analysis of highly polar pesticides in plant samples by LC-MS/MS and simultaneous extraction with methanol (QuPPe-PO-method, 2023). The procedure was as follows:

Samples of macerated weed were prepared to obtain a standard reference dilution curve. 5 g of each experimental sample and a blank matrix (grass without glyphosate) were homogenized and crushed. To each sample, 10 ml extraction buffer (1% acidified ethanol with formic acid) was added in a Falcon tube and vortexed for 5 min. Sonicated for 15 min, then frozen at −18°C for 90 min. Centrifuged at 6000 rpm for 10 minutes; 1 ml of supernatant was extracted from each sample and transferred to a vial using a sterile syringe with a 0.22 µm membrane. A 1:10 dilution of the filtered supernatant was prepared and 1 mL of the dilution was transferred to a plastic vial with a pre-punched cap to maintain the detection interval. Samples were transferred to a UPLC-MS/MS chamber for determination.

### Test organisms

*E. andrei* earthworms were provided by the Olmos company (San Marcos de León, Xico, Veracruz, Mexico). Four hundred sub-adult juvenile earthworms (without developed clitellum) were collected manually, with a similar biomass (0.2 ± 0.03 mg), and 4 kg of coffee pulp compost were also collected. Prior to the establishment of the experiments, in the laboratory of the Institute of Biotechnology and Applied Ecology (INBIOTECA for its initials in Spanish) of the University of Veracruz, the earthworms were kept in coffee pulp compost at ambient temperature with 80-90% humidity.

### Laboratory composting of *Pseudechinolaena polystachya*

For the composting of the weed contaminated with the different commercial doses of glyphosate (Low, Medium, High and Control) was placed in plastic boxes with 90% humidity for a period of 22 days, with a temperature of 55° C, 29° C and 25° C for 12, 5 and 5 days, respectively.

### Experimental design

The glyphosate toxicity essays were conducted in accordance with the International Standards for Avoidance Toxicology ISO17512-1 (ISO 2008) and Acute Test ISO11268-1 (ISO 2012).

Each experiment was set up under a completely randomized design, incorporating the four treatments applied in the field; the weeds collected from each treatment were divided into five replicates.

The experiment incorporated 50 juvenile earthworms (without developed clitellum) per treatment, with 10 earthworms per replicate. Each replicate received 150 g of compost at a ratio of 1:15 g. The duration of the avoidance bioassay (repellency and/or rejection) and the acute test was 48 hours and 14 days, respectively. At the conclusion of the bioassays, the earthworms were extracted from the compost, washed, and weighed.

### Avoidance essay

To perform the avoidance essay according to ISO-17512-1 (ISO, 2008), a completely randomized design was installed, plastic boxes (20×10×8 cm) were used; each box was divided into two equal sections (A and B) with plastic separator (10×8×0.5 cm) introduced vertically. In the first section (A) were placed 75 g of compost without glyphosate (Control) and in the second section (B) were placed the 75 g of compost contaminated with the respective glyphosate dose (Control, Low, Medium, High); then the separator sheet was removed; subsequently, in the indentation between the two sections (here they have the possibility to dig and select the habitat; section A and/or B), were introduced 10 juvenile earthworms of *E. andrei* with similar biomass (0.2 ± 0.03 mg) were introduced. Finally, the plastic boxes were covered and cultured in an incubator at 25 ± 2 °C and 80-90% humidity.

At 48 hours (end of the essay), in the plastic boxes of each of the treatments, between sections A and B, the separator sheet was introduced. In each section, live earthworms (responding to the mechanical stimulus) were collected, washed, dried and weighed. Missing earthworms were considered as dead and/or disintegrated during the essay.

### Acute Test Essay

The acute test ISO-11268-1 (ISO, 2012) was established based on a completely randomized design to evaluate *E. andrei* survival to exposure of weeds contaminated with four doses of glyphosate (ml/L). In plastic boxes (20×10×8 cm) as replicates, 150 g of compost (80-90% moisture) were placed with the respective treatments. Then 10 juvenile *E. andrei* earthworms with similar biomass (0.2 ± 0.03 mg) were introduced; subsequently, the plastic boxes were covered and reared in an incubator at 25 ± 2 °C, for 14 days.

At 7 and 14 days, in each of the replicates of the four treatments, earthworms were collected, washed, dried, weighed and counted. Missing earthworms were considered dead and/or disintegrated during the essay.

## Results and discussion

### Characterization of weeds

In the study area, the weeds collected in the coffee plantation correspond to a single species:

#### Pseudechinolaena polystachya

*P. polystachya* is a perennial weed species that grows in grasslands, shrublands and in cocoa and coffee plantations; it reaches about 50 cm in height (Mondragón et al., 2009). In the coffee plantation and its surroundings there are no records of the benefits or limitations it represents for coffee cultivation (Castro et al. 2019; Real-Luna et al. 2021). The average density recorded in the coffee plantation was 375 ind/m^2^ with an average productivity of 12 ± 10.8 g/m^2^ of dry matter. The nutritional quality of *P. polystachya* is distinguished by having 10.48% protein, 10.09% ash, 52.8% neutral detergent fiber, 27.7% acid detergent fiber and 4.5% acid detergent lignin (Mosisa 2021) and a Carbon/Nitrogen ratio of 44.94/3.19.

### Analytical determination of glyphosate

At 10 days after glyphosate application, the weeds of each treatment reflected the effect of the herbicide expressed as a change in coloration from green to yellowish. The results of the analytical determination showed that the glyphosate applied (Control, Low, Medium and High) to the weed *P. polystachya* was equivalent to 171.75 ± 0.00, 540.27 ± 19.59, 993.05 ± 80.27 and 1172.04 ± 2.19 mg/kg, respectively, with an average absorption of 0.0, 259.32, 437.03 and 398.5 mg/kg; that is, 0, 48, 44 and 34% of the applied glyphosate.

During composting (22 days), it was observed that the foliage of the weed *P. polystachya* changed in flaccidity (from minor to major) and coloration (light to dark green) as the glyphosate dose increased (Control, Low, Medium and High). Glyphosate favors decomposition, as documented in previous studies (Gaupp-Bergahusen et al. 2015; Vivanco et al. 2023) and associated with the activity of microorganisms (Lorch et al., 2021; Raffa et al., 2021; Patison et al., 2024).

### Avoidance essay

Some earthworm species do not avoid glyphosate-contaminated sites and are efficient mediators of herbicide degradation (Owagboriaye et al., 2020b; Dada et al., 2021; Zaller et al., 2021). At the end of the avoidance test (48 h) 99.5% of *E. andrei* earthworms were found alive (n=200).

The variation in biomass loss of *E. andrei* during the avoidance test was not significant among glyphosate treatments (F = 1.16, p = 0.33). In Figure 1 it can be observed that the trend in earthworm biomass loss was low but it is observed that it was higher in the treatments with half dose (1.92 ± 0.088 g) and Low dose (1.60 ± 1.55 g), respectively; while in the Control (1.83 ± 0.025) and high (1.73 ± 0.28 g) treatments had an intermediate loss. Previous studies have documented that glyphosate increases the incorporation of organic matter (more digestible and labile nutrients) into the soil; for example, it has been recorded that the presence of the herbicide was associated with an increase in earthworm biomass (Zaller et al., 2014; Gaupp-Bergahusen et al., 2015) and increased bacterial activity (Stellin et al., 2018; Machado and Kirel et al., 2021; Patison et al., 2024).

**Figure 1.**
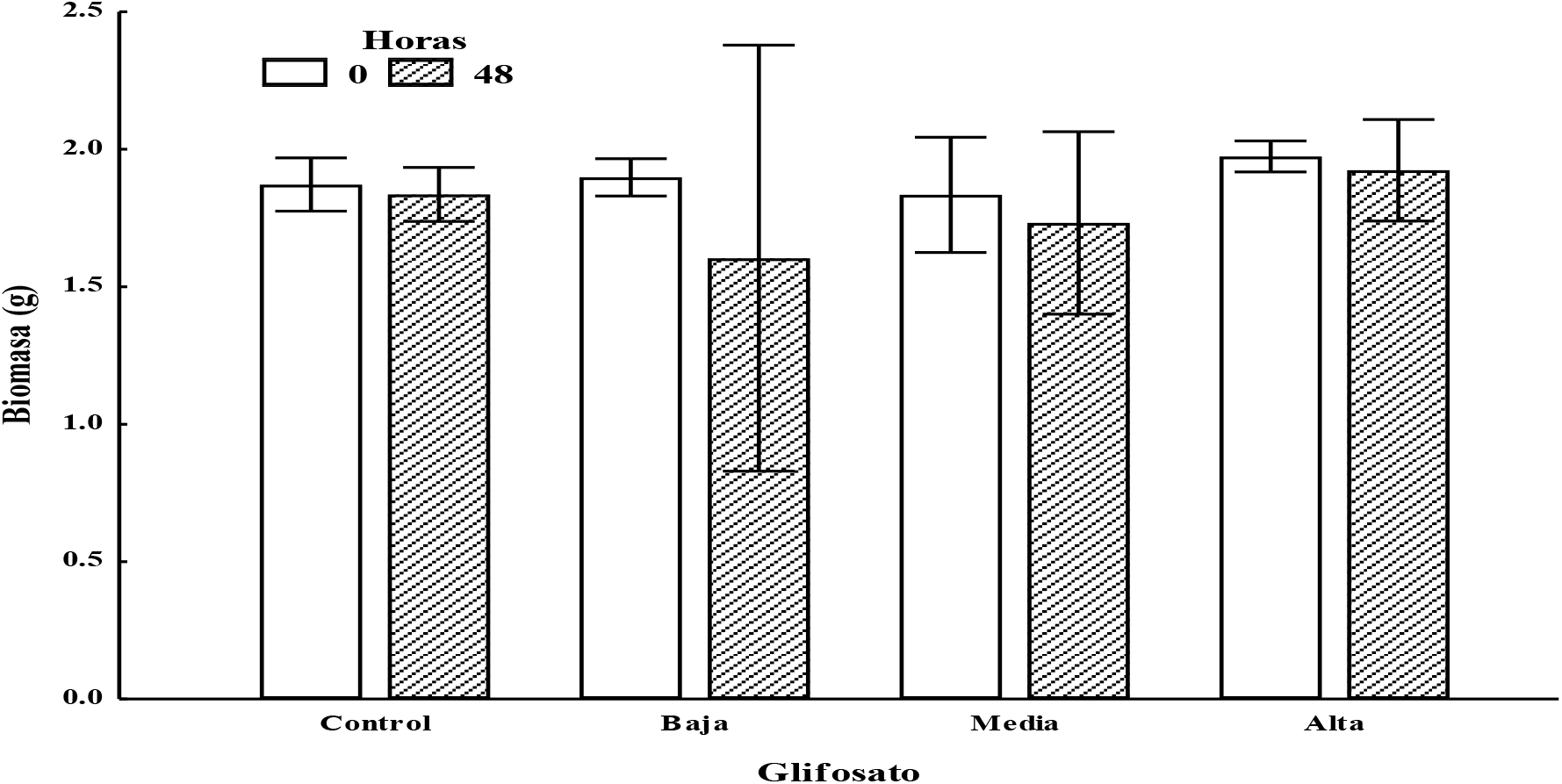
Initial and final biomass of earthworm Eisenia andrei after 48 hours of exposure of compost of the weeds Pseudechinolaena polystachya contaminated with different doses of the herbicide glyphosate. Values are means and vertical lines represent standard error (n = 10).

On the other hand, results at 48 hours showed that the avoidance response to glyphosate exposure varied significantly among treatments (F = 2.46, p = 0.09). Earthworms did not reject or avoid (>80%) compost of the weed *P. polystachya* contaminated with different doses of glyphosate. *P. polystachya* compost is a habitat that provides food, shelter, etc. (ISO 2008; Pochron et al., 2021). The Control, Low, Medium and High treatments had E. andrei repellency of 44 ± 37.8, 30 ± 42.4, 8.0 ± 8.4 and null, respectively (Figure 2). These results are contrary to what was expected and recorded in previous studies (Casabé et al., 2007; García-Pérez et al., 2014; Stellin et al., 2018). Van Hoesel et al., (2017) and Zaller et al., (2018) reported that glyphosate is not associated with weed foliage decomposition. However, in this study, it was observed that contaminated *P. polystachya* weed compost disintegrated more with increasing glyphosate concentration as recorded by Vivanco et al., (2023); for example, in the Alta treatment, no earthworm repelled the compost (Hernández et al., 2017), while in the Control treatment (without glyphosate) the rejection was higher (ISO 208; Pochron et al., 2021). The results suggest that *E. andrei* accepts *P. polystachya* compost with glyphosate as food as it is more digestible and nutritious (Gunadi and Edwards 2003; Rasche 2004; Gaupp-Bergahusen et al., 2015).

**Figure 2.**
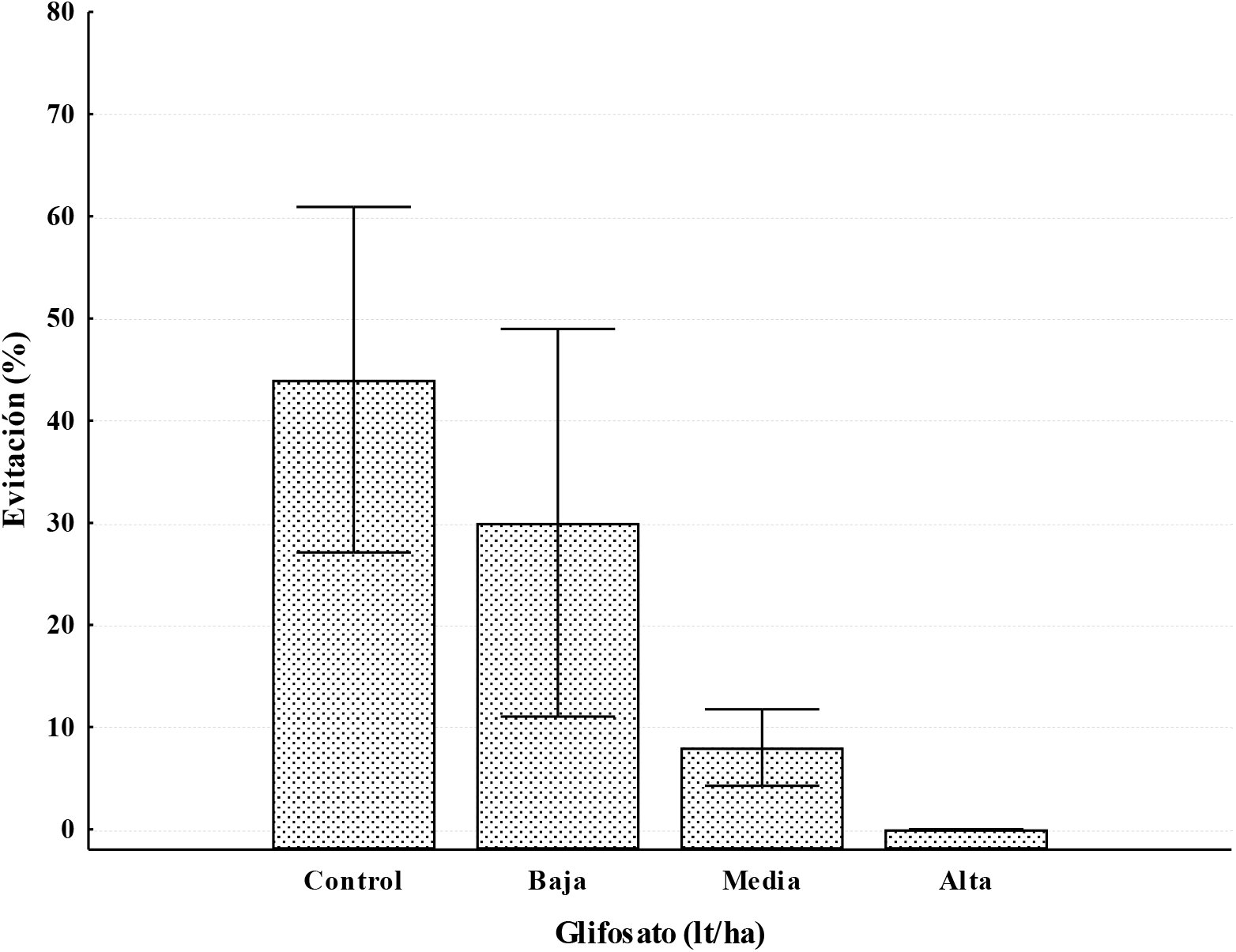
Avoidance behavior of the epigean earthworm Eisenia andrei after 48 hours of exposure to compost of the weeds Pseudechinolaena polystachya contaminated with different doses of the herbicide glyphosate. Values are means and vertical lines represent standard error (n = 10).

### Acute exposure essay

#### Earthworm biomass

Increased earthworm biomass is associated with the quality and quantity of nutrients available in organic matter and soil (Gunadi and Edwards 2003; Rasche et al., 2004; Stellin et al., 2018) while biomass loss has been associated with a toxicological response (Correrira and Moreira, 2010; García-Pérez et al., 2014; Pochron et al., 2018) and a detoxification process of earthworms (Lescano et al., 2020; Owagboriaye et al., 2020a; Pochron et al., 2021). The results of the present study showed that, at 7 and 14 days, among glyphosate-contaminated *P. polystachya* composts (Control, Low, Medium and High), the biomass of *E. andrei* varied significantly (F = 2.92, p = 0.04).

In contrast to what was expected, at 7 days the earthworms in the treatments with Medium and High doses registered a biomass gain that was associated to the fact that the weed was more composted and/or in decomposition (greater digestibility and availability of nutrients); while in the Control and Low treatments the earthworms lost biomass probably due to less decomposition (less digestible) of the *P. polystachya* compost (Figure 3). However, at 14 days, in all treatments *E. andrei* lost biomass, highlighting the High and Medium treatments (higher concentration of glyphosate) for losing less biomass. Several studies have reported epithelial alterations (Giunta and Jauregui, 2013) and modifications to the intestinal bacterial colonies of earthworms (Ceceña et al., 2022; Narváez et al., 2022; Want et al., 2023).

**Figure 3.**
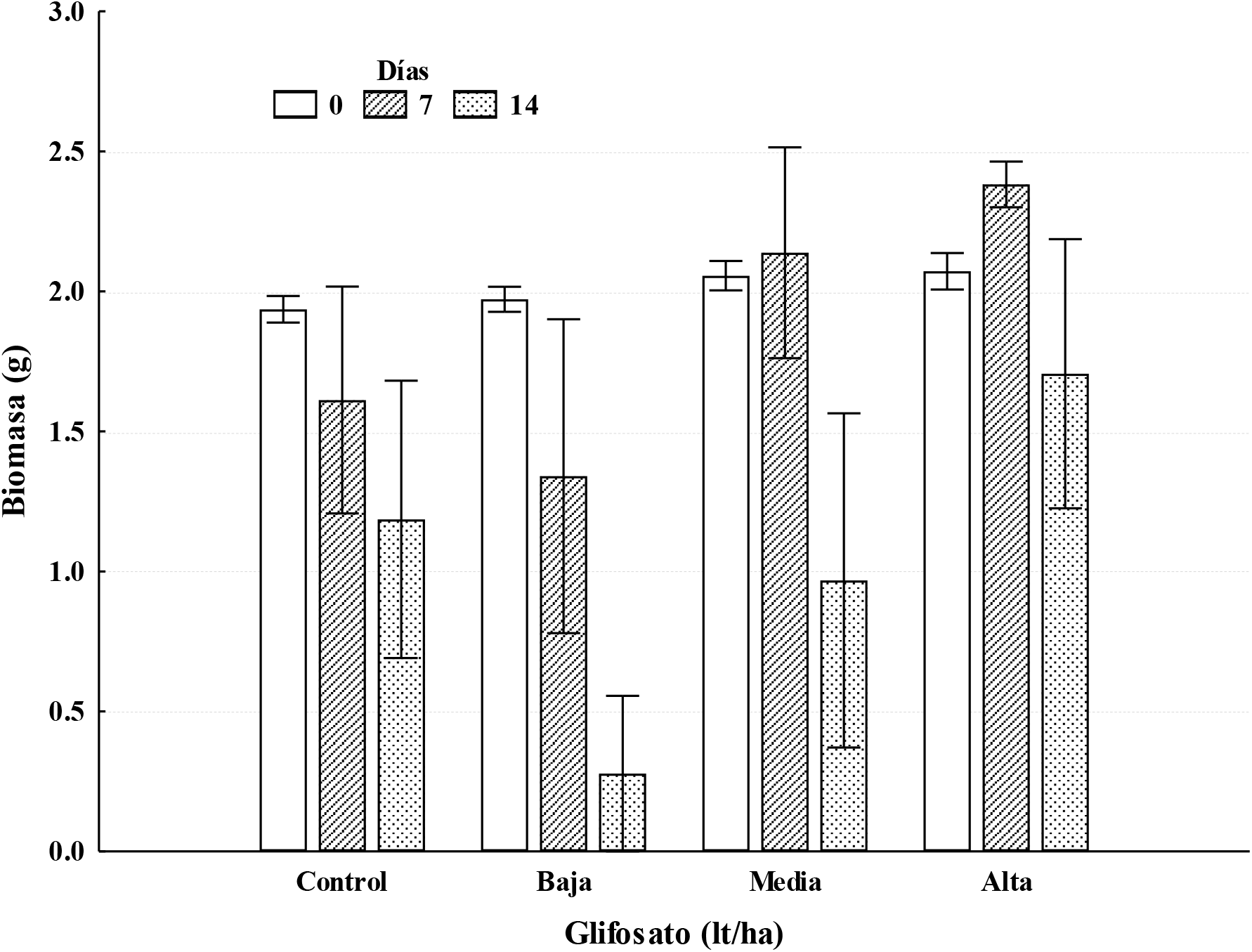
Initial and final biomass of the epigean earthworm Eisenia andrei after 7 and 14 days of exposure to compost from the weeds Pseudechinolaena polystachya contaminated with different doses of the herbicide glyphosate. Values are means and vertical lines represent standard error (n = 10).

#### Survival percentage

The response of earthworms to glyphosate varies among species (Doublet et al., 2009; Zaller et al., 2018; Lorch et al., 2021). Some species are more tolerant to glyphosate without causing death (Andrea et al., 2004; Maitre et al., 2010; Pochron et al., 2021); for example, *Eudrilus eugenianae* and *Libyodrilus violaceous* reduce soil glyphosate concentrations (Owagboriyae et al., 2020a; Palafox et al., 2021). The results of our study showed that the variation of *E. andrei* mortality among treatments was not significant at 7 and 14 days (F = 0.88, p = 0.45). It has been documented that *E. andrei* can survive a few days without feeding (Mósqueda et al., 2019), however, authors such as Garcia-Perez et al., (2020), Yang et al., (2022) and Silva and Pelosi (2024) have reported biomass loss and developmental delay in earthworms. At the end of the experiment (14 days) the highest average mortality of earthworms was registered for the Low treatment (86 ± 31.3%) and the lowest in the High treatment (28 ± 43.8%). The Medium treatment had a mortality of 60 ± 77.2%, while the Control treatment had a mortality of 40 ± 54.8% (Figure 4).These results suggest that at 7 and 14 days *E. andrei* mortality is inversely associated with compost nutrient availability influenced by glyphosate concentration in *P. polystachya*; that is, mortality was higher (Low and Medium treatments) and lower (Control and High treatments) in compost with low and high nutrient availability and glyphosate concentration, respectively (Pochron et al., 2018; Owagboriaye et al., 2020b; Zaller et al., 2021). In contrast at 14 days, mortality was due to an effect of glyphosate, because in the control treatment earthworm death was constant while in the glyphosate treatments mortality after 7 days increased markedly at the end of the experiment (Correira and Moreira, 2010; Garcia-Perez et al., 2014; Zaller et al., 2021; Yang et al., 2022).

**Figure 4.**
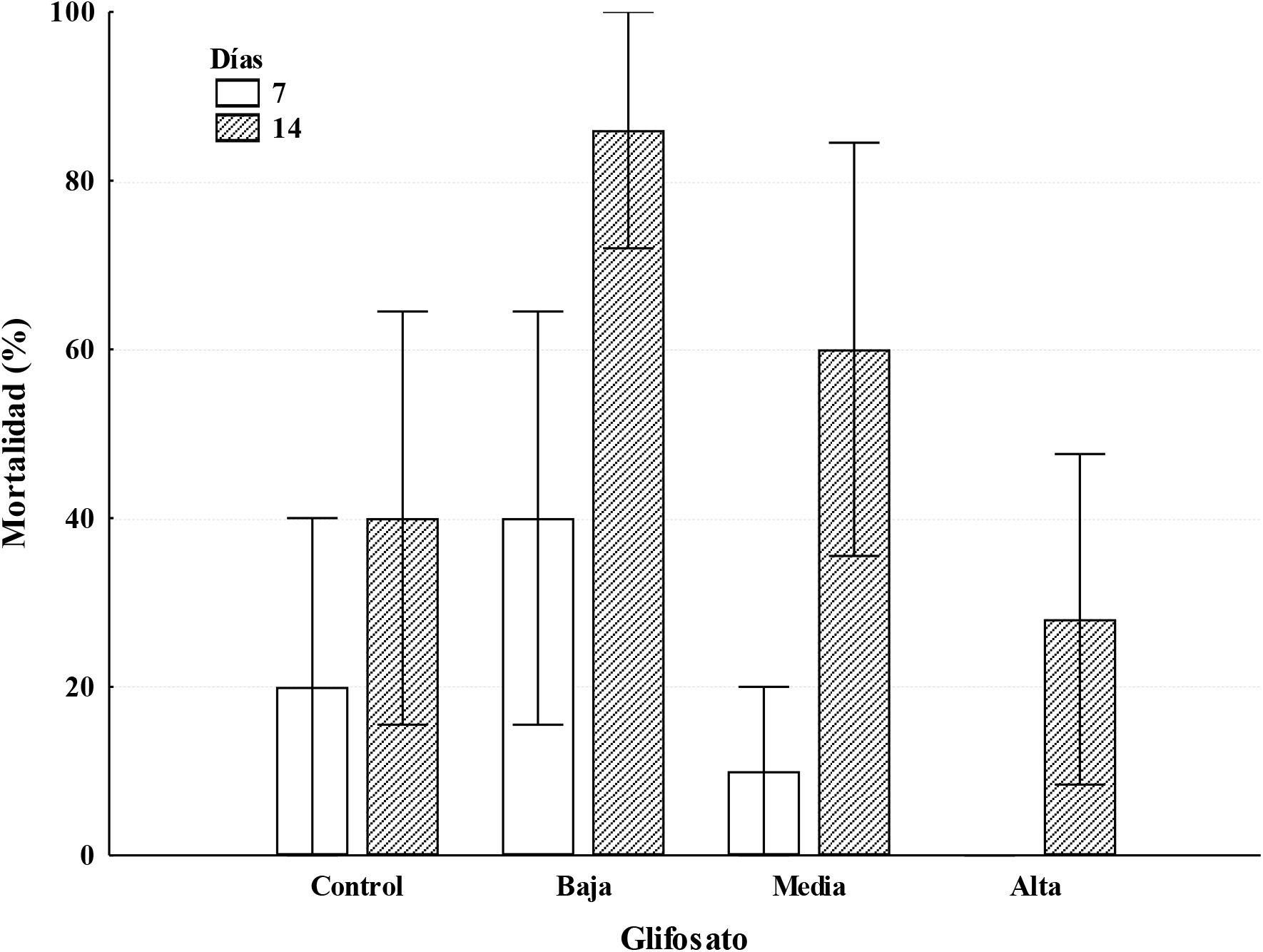
Mortality of the epigean earthworm Eisenia andrei after 7 and 14 days of exposure to different weeds (Pseudechinolaena polystachya) composts contaminated with the herbicide glyphosate. Values are means and vertical lines represent standard error (n = 10).

## Conclusion

The results indicated that in the shaded coffee plantation the dominant weed was the species *P. polystachya*. The application of glyphosate herbicide at commercial doses of 2 to 6 l/ha concentrated from 259.3 to 398.5 mg/kg of dry matter. The avoidance test was less sensitive than the acute test; that is, *E. andrei* did not repel glyphosate-contaminated weed composts. However, the acute test showed that increasing the herbicide concentration reduced biomass loss and mortality of *E. andrei* due to increased compost decomposition. The study reveals the toxicity of the recommended dose of glyphosate on the survival of *E. andrei*.

## Acknowledgments

The authors thank the Postgraduate Program for the Master of Science in Agricultural Sciences of the Veracruz University.

